# Macroevolutionary changes in gene expression response to an immune stimulus across the diversity of fishes

**DOI:** 10.1101/2024.12.13.628417

**Authors:** Ben A. Flanagan, Lauren E. Fuess, Milan Vrtílek, Andrea Roth-Monzón, Daniel I. Bolnick

## Abstract

Our understanding of the vertebrate immune system is dominated by a few model organisms such as mice. This use of a few model systems is reasonable if major features of the immune systems evolve slowly and are conserved across most vertebrates, but may be problematic if there is substantial macroevolutionary change in immune responses. Here, we present a test of the macroevolutionary stability, across 15 species of jawed fishes, of the transcriptomic response to a standardized immune challenge. Intraperitoneal injection of an immune adjuvant (alum) induces a fibrosis response in nearly all jawed fishes, which in some species contributes to anti-helminth resistance. Despite this conserved phenotypic response, the underlying transcriptomic response is highly inconsistent across species. Although many gene orthogroups exhibit differential expression between saline versus alum-injected fish in at least one species, few orthogroups exhibit consistent differential expression across species. This result suggests that although the phenotypic response to alum (fibrosis) is highly conserved, the underlying gene regulatory architecture is very flexible and cannot readily be extrapolated from any one species to fishes (or vertebrates) more broadly. The vertebrate immune response is remarkably changeable over macroevolutionary time, requiring a diversity of model organisms to describe effectively.

## Introduction

Advancements in immunology have largely been driven by research on a few model organisms including mice, macaques, and zebrafish. While these models have been transformative in our understanding of the immune system, their application to humans relies on the evolutionary stability of the immune process across tens to hundreds of million years of evolutionary divergence. Emerging evidence suggests that the structure and function of the vertebrate immune system varies greatly, at several scales of biological organization. Immune variation can be observed among individuals within a given population. For example, severity of SARS-CoV2 is genotype dependent; individuals homozygous for the *e4* allele at *ApoE* gene display increased mortality (Kuo et al. 2020). This genetic variation within populations can then be a target for natural selection, enabling rapid evolutionary change within populations through time, leading to genetic differences between populations and thus species. For instance, the frequency of alleles at Major Histocompatibility Complex (MHC) loci diverges between animal populations subjected to different parasite communities (Wegner et al. 2003; O’Connor et al. 2018; Osborne et al. 2017). These differences have been thought to contribute to the evolution of reproductive isolation and speciation. As a result, immune genotypes and functions can differ appreciably between even closely related species. Given this documented rapid evolution of immunity within and between closely related species, it seems improbable that immune function should be highly conserved across deeper evolutionary time. At a minimum, we should seek to understand what features of immunity are conserved, and what features are divergent across animals (e.g., Gilbertson & Weinmann 2021).

Major evolutionary transitions in the immune system certainly occur within vertebrates. Within fishes, some supposedly essential features of adaptive immunity have turned out to be surprisingly dispensable. MHC emerged roughly 500 million years ago at the origin of chordates, and are one of the most studied immune genes. MHC is present in nearly all vertebrates, appearing as a fundamental and canonical aspect of adaptive immune function. Yet, there is substantial turnover in the number and identity of MHC paralogs between closely related primates (Lyn Fortier & Pritchard 2024). Sometimes whole categories of MHC are lost, such as the non-functionalization of MHC Class II in atlantic cod (*Gadus morhua*) (Star et al. 2011) and its close relatives (Malmstrøm et al. 2016). Even more dramatically, angler fish have lost many components of their adaptive immune system, enabling an evolutionary transition to extreme size sexual dimorphism (Brownstein et al. 2024) and male reproductive parasitism (Swann et al. 2020; Dubin et al. 2019). It is essential to understand whether these examples of immune function loss (or change) are exceptions, restricted to a few lineages, or are typical of other immune functions and gene families.

Such changes in immune function may prove to be common across vertebrates, but there have been few systematic evaluations of immune function across a diverse swath of vertebrate species. Comparative immunology has been limited in part by the difficulty of studying immune responses in many species simultaneously, especially considering the need for species-specific reagents such as monoclonal antibodies to measure many immune traits. And yet, the advent of genome and transcriptome sequencing has facilitated high throughput comparisons of gain or loss of genes across many species’ genomes, or changes in transcriptional responses. Comparative transcriptomics is emerging as an especially promising tool for comparative immunology (Chen et al. 2023; Yang et al. 2022), without the need to develop species-specific molecular probes. One can measure differential expression of genes within each of many species, subjected to a standardized immune challenge. We can then compare differential expression among many diverse species, to ask: is the vertebrate immune response highly conserved across most vertebrates? Or, is the immune response evolving so quickly that each species responds to a given immune challenge with a unique set of transcriptional changes? Almost certainly, the answer lies somewhere between these extremes, and comparative transcriptomics can let us quantify the extent to which immune responses have a shared, or unique, transcriptional basis across a wide range of animals.

Here, we use comparative transcriptomics to test whether a sample of fish species from across the full span of the teleost tree of life exhibit a similar response to a standardized immune challenge. Many species of fish (and other vertebrates) develop extensive fibrosis during an inflammatory immune response (Vrtílek & Bolnick 2021). Fibrosis involves the formation of an extensive extracellular matrix of protein fibers (collagen, fibrinogen), produced by an evolutionarily ancient cell type (fibroblasts). Fibrosis is essential in wound healing, but is also a common feature of inflammation (Wynn & Ramalingam 2012). This fibrosis response can be beneficial: in some fish species (e.g., threespine stickleback), fibrosis has been found to confer protection against intraperitoneal tapeworm infection (Lohman et al. 2017; Weber et al. 2021). Fibrosis is also widely considered to be a pathological side-effect of inflammation, and is involved in a variety of human diseases (Henderson et al. 2020). In stickleback, the immune benefits of fibrosis are countered by reduced fecundity (De Lisle & Bolnick 2021) and reduced mobility (Matthews et al. 2023). Consequently, fibrosis responses evolve rapidly within, and diverge between, stickleback populations (Hund et al. 2022), which exhibit microevolutionary changes in fibroblast activation genes (Fuess et al. 2021).

In stark contrast to this rapid evolution of fibrosis in stickleback, another study found that fibrosis responses are highly conserved across teleosts (Vrtílek & Bolnick 2021). Intraperitoneal injection of an immune adjuvant (alum, widely used in vaccines to stimulate inflammation) causes a strong fibrosis response within the body cavity of stickleback (Hund et al. 2022). Extending this assay to 17 species of fish sampled from across the teleost phylogeny, Vrtilek and Bolnick (2021) found that nearly all species initiate fibrosis after alum exposure. This result reveals a puzzling contrast: fibrosis is both highly conserved among teleost species spanning over a hundred million years (Vrtílek & Bolnick 2021), but diverges between stickleback populations separated for a mere 10,000 years (Weber et al. 2022). This contrast is an example of the broader contradiction of rapid immune evolution, and conservation of immune features, in vertebrates. Usefully, we can apply a standardized immune challenge (alum), which elicits a standard phenotypic response (fibrosis) in most fish species. Here, we pose the question: is this similar phenotypic response to alum generated by a standard transcriptomic response, shared across diverse teleosts? Or, do different fish species respond to alum in unique ways, indicating significant changes in the transcriptomic pathways responding to an immune challenge and driving fibrosis?

To answer these questions, we generated reference transcriptomes for 14 species from the Vrtilek and Bolnick (2021) study. We sequenced the head kidney (pronephros) transcriptomes of the experimental fish that had been injected with either saline (controls) or alum solution, using head kidneys archived from the Vrtilek and Bolnick experiment. Within each species, we test whether each transcript family (orthogroup) exhibits differential expression in response to alum injection (relative to saline controls). Then comparing across species, we test whether this differential response is shared among species, or varies among species. That is, do we observe a treatment*species interaction effect in which an orthogroup is up- or down-regulated in response to alum in some species, but not others (or, responds in opposite directions depending on the species). We then examine the most differentially expressed genes across any teleosts, to identify what biological functions are responding to alum. For instance, do we see consistent overrepresentation of fibrosis-related or inflammation-related genes, even if the specific orthogroups are inconsistent from species to species? Are the changes in expression, among species, associated with the extent to which species develop fibrosis after alum injection? We show, below, that each species exhibits a transcriptional response to alum (even the few species that do not respond with fibrosis). But, this transcriptional response is unique to each species, at the level of transcript orthogroups. However, at the level of gene function (e.g., gene ontology category), we see some weakly similar responses across species separated by hundreds of millions of years of evolutionary divergence.

## Methods

### Fish selection and experimental methods

To gain an understanding of the evolution of gene expression associated with the formation of peritoneal fibrosis across ray-finned fishes (Actinopterygii), we intentionally selected species spread across the phylogenetic diversity of Actinopterygii which were small and commercially available. The 17 fish species were housed in laboratory aquaria and underwent experimental injections as described in (Vrtílek & Bolnick 2021). Because tapeworm proteins only limitedly induces fibrosis in ray-finned fishes (Vrtílek & Bolnick 2021), we focus on the gene expression response to alum, an immune adjuvant which activates the innate immune response (Kool et al. 2012) and broadly induces peritoneal fibrosis in Actinopterygii (Vrtílek & Bolnick 2021). Fish were peritoneally injected through their left flank with volume-mass corrected solutions at 20 uL per 1g of average weight for a given species and the species included 1x phosphate-buffered saline (PBS), and a 1% alum solution. These fish were euthanized five days following injection, but because stickleback and trout were raised at a lower temperature, they were euthanized ten days post injection. Following euthanasia, each sample was scored ordinally for fibrosis which ranged between 0 - 3 where 0 indicates no fibrosis with freely moving internal organs and no fibrotic attachment to the peritoneal wall. Level 1 fibrosis is when internal organs adhere together and move as a unit. Level 2 fibrosis occurs when organs attach to the peritoneal wall but can be detached while fibrosis level 3 was the most extreme fibrosis phenotype where organ adhesion to the peritoneal wall could not be reattached without the peritoneal lining tearing apart and remaining on the internal organs when forcibly opened. This ordinal fibrosis score is described in (Vrtílek & Bolnick 2021) and is similar to the ordinal classification of fibrosis presented in (Hund et al. 2022). The ordinal scoring is highly repeatable among independent observers (Bolnick et al. 2024).

In addition to scoring fibrosis, the head-kidney tissue was isolated and stored in RNAlater. We focused on characterizing the specific head-kidkney transcriptome response because in teleosts it is one of the tissues responsible for initializing the immune response, both innate and adaptive (Zapata et al. 2006). Of the 17 species used in the Vrtilek and Bolnick experiment, here we study 14 species whose head kidneys yielded sufficient quantities of high quality RNA, and which had (or we could generate) reference transcriptomes from head kidneys.

### Reference transcriptome sequencing

We extracted RNA from frozen head kidney samples using MagMax96 Total RNA Isolation kit, following manufacturer’s instructions. RNA yield was quantified using a Ribogreen fluorescent stain compared with a standard curve, with fluorescence measured on a Magellan 96-well plate fluorometer. While some species had sequences available from head kidney tissues (Table S1), there were species which lacked available head kidney sequences, so we generated *de novo* references using paired end 150bp sequencing through Novogene (20 million reads per sample). We sequenced and assembled *de novo* transcriptomes for *Polypterus senegalus, Hyphessobrycon erythrostigma, Tateurndina ocellicauda*, *Sphaeramia nematoptera, Chromis viridis, Salarias fasciatus, Nothobranchius furzeri, Lepomis macrochirus, Dichotomyctere nigroviridis,* and *Helostoma temminckii.* We focused on head kidney transcriptomes because we wished to build references appropriate to the transcriptomes we later examine with TagSeq, using the same tissues.

### Transcriptome assembly and comparative transcriptomics

While some of the fish species had previously published transcriptomes, we wanted to ensure congruent assembly methods occurred for all species, so we independently *de novo* assembled transcriptomes for all species using previously published data where available, and conducting full-length RNA-seq for a subset of species which had no head kidney RNA-seq data available (Table S2). *De novo* head kidney transcript assembly using previously published sequences included *Danio rerio (Pasquier et al. 2016), Gasterosteus aculeatus (Huang et al. 2016), Oncorhynchus mykiss (Sun et al. 2023),* and *Oreochromis niloticus (Hou et al. 2023).* For each species including those sequenced here and species with previously published data, we *de novo* assembled the head kidney specific transcriptome using rnaSPAdes with paired end reads (Bushmanova et al. 2019). Then, to identify coding regions of the transcriptome we ran *Transdecoder (Haas et al. 2013)* which first identifies the longest open reading frame with a minimum length of 100aa and then searches for protein domains with similarity to the Pfam-A Pfam database using *HMMER3* (hmmer.org). Lastly, *Transdecoder* predicts candidate coding regions in the transcriptome.

The resultant *Transdecoder* (Haas et al. 2013) predicted peptide sequences for all species were used to identify orthologs, orthogroups, and infer a species tree based on duplication events through *Orthofider2* (Emms & Kelly 2019; Emms & Kelly 2015) using *DIAMOND* (Buchfink et al. 2015) and the *STRIDE* (Emms & Kelly 2017) method to infer species tree. Previously, studies attempting to characterize the evolution of gene expression were limited by focusing on genes which were single-copy in all species. Moving beyond single copy orthogroups to include orthogroup-level comparisons with species transcript content variability, we begin to understand how gene families with shared, potentially constrained function, evolve across phylogenetic diversity.

### 3’TagSeq for differential expression of experimental fish

For samples from the injection experiment, we extracted RNA from head kidneys of saline injected and alum injected individuals of each species (sample sizes in Table S2). We used ribogreen to quantify and normalize the extracted RNA to 20ng/ul, and sent these to the Genome Sequencing and Analysis Facility at the University of Texas at Austin for 3’TagSeq, a cost-effective approach that sequences just the 3’ ends, providing cheaper and more precise estimates of relative expression levels (albeit without information about splicing variants). 3’TagSeq libraries were constructed according to (Lohman et al. 2016; Lohman et al. 2017) based on methods presented in (Meyer et al. 2011)), aiming for 5 million reads on average per sample.

To generate read counts per transcript from the *de novo* head kidney transcriptomes assemblies, we used the coding sequence output from the *Transdecoder* predict algorithm. Using the pseudo-mapper Salmon (Patro et al. 2017) which maps and counts in a single step, we first indexed the transcriptome and subsequently quantified reads mapping to each transcript. After quantifying reads for each transcript, counts were imported into *R (R Core Team 2024)* and were normalized using the mean-of-medians approach described in *DEseq2* (Love et al. 2014).

### Orthogroup differential expression

For transcripts with a shared evolutionary history which were placed into orthogroups, the normalized transcript counts were summed for all transcripts which have membership in a given orthogroup for each sample individually. The resultant orthgroup count matrix was then analyzed using *DEseq2* (Love et al. 2014) to estimate orthogroup differential expression.

### Statistical analysis

Orthogroup expression was summed for all transcript members of a given orthogroup for each species, then to determine the treatment effect for each orthogroup and species, we initially fit a linear model in *R* (lm). Then, we estimated the interaction effect for treatment and species for each orthogroup then subsequently estimated the effect size (η2) for treatment and the interaction effect which allows us to determine whether species-specific treatment effects (treatment by species interaction) or shared treatment effects dominate variance in orthogroup expression.

To determine if differential orthogroup expression response to the alum treatment was consistent across species, for each orthogroup and species, we estimated the Wald test statistic using *DeSeq2* (Love et al. 2014). Then to check whether the overall transcriptome-wide response was similar between species, we calculated correlations between all pairs of species, comparing the vectors of orthogroup Wald test statistics from each species.

Similarly, to determine if gene ontology (GO) enrichment was consistent across all species, we first determined which transcripts were differentially expressed between treatments, within each species, using *DeSeq2* (p-value < 0.1) *(Love et al. 2014)*. Then, for each *de novo* transcriptome assembly, we determine GO membership for each transcript using the web application TRAPID 2.0 (Bucchini et al. 2021). We then estimated the enrichment of differentially expressed transcripts for each GO term using a Fisher’s exact test in *R* (fisher.test) estimated by an odds ratio test statistic. The resultant odds ratios for all GO terms were pairwise correlated for all species.

### Data and code availability

Sequences for the de novo reference transcriptomes, assembled reference transcriptome, and 3’TagSeq data are archived on NCBI (to be done upon submission). The 3’TagSeq read count matrices, orthogroup assignments, and metadata for experimentally injected fish, and supporting code are archived publicly on Dryad (to be done upon submission).

## Results

Sequencing and *de novo* assembling head kidney specific transcriptomes from 14 diverse ray-finned fish species resulted in 65-92% of the assembled transcripts being assigned to an orthogroup (Figure 1A). The comparative approach identified only 20 single-copy orthologs across all the 14 species. In contrast, 7,528 orthogroups had at least one transcript from all species, out of 73,286 total orthogroups identified, with a mean orthogroup size of 10.8 distinct transcripts. This paucity of single-copy orthologs is unsurprising given that our phylogenetic sampling spans at least two major whole genome duplication events. The head kidney *de novo* assemblies were used to infer the phylogenetic relationship among the experimental species and it is congruent with the current ray-finned fishes phylogeny (Hughes et al. 2018) (Figure 1B).

**Figure 1:**
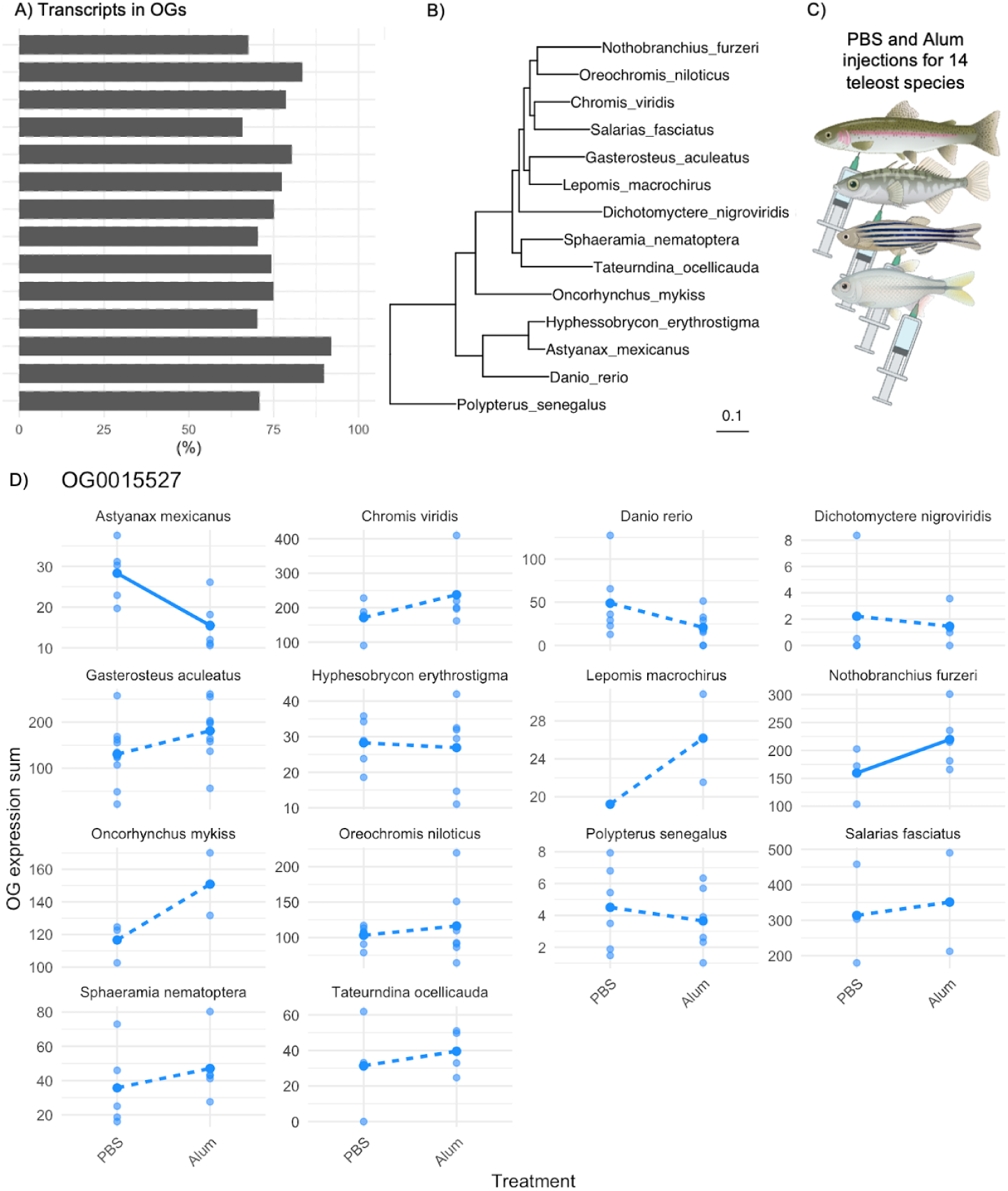
A) Percentage of transcripts from each species assigned to orthogroups (OG). B) A species tree inferred from all genes (STAG) with root inferred using duplication events (STRIDE) with branch lengths equivalent to the average number of substitutions per site across gene families. C) Experimental diagram illustrating intraperitoneal injections for the PBS (control) and an Alum stimulus (immune adjuvant) for four representative species (top to bottom: *O,mykiss*, *G. aculeatus*, *Danio rerio*, and *A. mexicanus*). D) Summed orthogroup (OG) normalized expression for a single-copy ortholog (OG0015527) using DESeq2’s median of ratios method. Each point represents treatment-specific OG summed expression for each sample within each species. where solid lines represent a significant treatment effect (p-value < 0.1) and dotted lines represent non-significant treatment effect (p-value > 0.1). Created with BioRender.com.

For many orthogroups, expression level varied among the species in our samples (96.76% of orthogroups show a main effect of species, p-value < 0.1). For example, differential expression in response to alum injection for orthogroup OG0007364 was negative in *S. nematoptera* (log2 fold change [lfc] of −1.63) but expression increased in *D. rerio* (lfc = 0.29). Blastp results indicate OG0007364 transcript members are derived from Target of ERG1 protein 1, *toe1*, which inhibits cell growth rates and induced TGF-beta expression (De Belle et al. 2003). In contrast, few orthogroups consistently respond to the alum injection treatment (20.34% of orthogroups had a significant main effect of alum treatment, p-value < 0.1). As an example, OG0011385 lfc (comparing alum injected vs control fish in each species) ranged from −0.98 (in *S. fasciatus)* to −0.05 (in *A. mexicanus*) indicating a moderately consistent down-regulation of this gene in alum-injected fish, though the magnitude of this change varied among species. The top blastp result for OG0011385 is *sap18*, a component of the SIN3-repressing complex which can direct the formation of a repressive complex to core histone proteins (Singh et al. 2010). The number of distinct transcripts assigned to orthogroups (e.g., gene family size) varied among species, and there was a weak negative correlation between the absolute value of lfc in gene expression (alum vs saline) and the number of transcripts assigned to an orthogroup (Pearson’s product-moment correlation = −0.04, t = −11.51, p-value > 2.2e-16). This indicates copy number minimally influences the estimated differential expression response to injection, based on summing the expression for all transcripts present in a species for a given orthogroup.

Instead of a uniform response, we observe species-specific transcriptomic responses to alum. In estimating the effect size of treatment and the treatment*species interaction by fitting a linear model (summed expression ∼ treatment + treatment*species) for each orthogroup, the interaction effect sizes are consistently larger than the treatment effect sizes (Figure 2A). Even for the orthogroup with the highest amount of variation explained by treatment (OG0000016, hemoglobin subunit alpha), treatment only explained 9.4% of the variation while the interaction of treatment and species explained a larger proportion of variation (19.3%) illustrated by the species-dependent slopes in response to alum injection (Figure 2C&D). Therefore, the response to alum is highly heterogeneous among fish species indicating there is no single gene regulatory program responding to alum consistently across all species. Similarly, there is no single gene regulatory program associated with high versus low fibrosis within species (Supplementary Figure S2).

**Figure 2.**
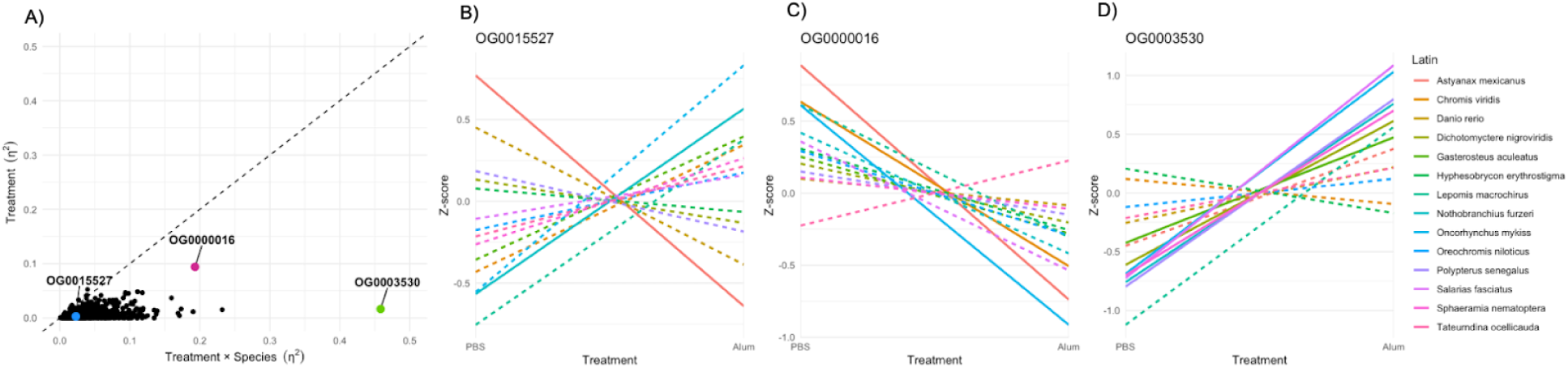
A) For each orthogroup summed normalized gene expression, we used a linear model to estimate the effect size (η^2^) for the effects of injection treatment (PBS versus Alum), and the treatment × species interaction on variation in the focal trait. Treatment η^2^ indicates the extent to which any orthogroup summed normalized expression diverges predictably from PBS to Alum. The η^2^ for the treatment × species interaction measures the extent to which treatment effect depends on the species for every orthogroup. The dashed line is a 1:1 line, for ease of visualization; points falling above this line have a larger treatment effect than interaction effect. Orthogroup expression for PBS and Alum injections across all species where solid lines represent a significant treatment effect (p-value < 0.1) and dotted line represents non-significant treatment effect (p-value > 0.1). B-D) Standard normal (z-score) expression for B) single copy ortholog (OG0015527), C) orthogroup with highest effect size of treatment (OG0000016), and D) orthogroup with highest effect size of the interaction of treatment and species (OG0003530)

Despite the lack of conservation at single orthogroups, there are some transcriptome-wide similarities between species’ responses to alum. On average across the transcriptome, orthgroups which are up- (or down-) regulated in one species, tend to be up- or down-regulated in other species. Specifically, there is a correlation between log2 fold changes associated with alum, when comparing pairs of species (Figure 3A&B). For example, the complete vector of log2 fold changes in *Tateurndina ocellicauda* and the corresponding lfc vector for *Oncorhynchus mykiss* are positively correlated (Pearson’s correlation coefficient = 0.12, t = 5.16, p-value = 2.71e-07) (Figure 3A). For all species-pairwise correlations, the distribution falls greater than zero (t-test, t = 9.44, p-value = 4.23e-15) (Figure 3B).. This correlation between orthogroup differential expression varies in strength depending on the pair of species being examined (Figure 3B). Some species pairs are more strongly correlated, implying more similar transcriptome-wide response to alum, than other pairs of species whose response is uncorrelated (*O. niloticus* and *T. ocellicauda*, Pearson’s correlation coefficient = 6.94e-05, t = 0.004, p-value = 0.99), or negatively correlated (Pearson’s correlation coefficient = −0.1, t = −4.49, p-value = 7.45e-06). Yet, overall there is a tendency for a similar general response to alum among species, though this is not observed for individual orthogroups. Further, we might expect species which are more closely related to show similar patterns of orthogroup differential expression, yet there is discordance between the hierarchical grouping of differential orthogroup expression and the inferred species tree (Figure 4). The evolution of differential orthogroup expression does not follow species diversification wherein species changes occur independently and may be contained to specific orthogroups with accelerated expression changes (Figures 3&4).

**Figure 3.**
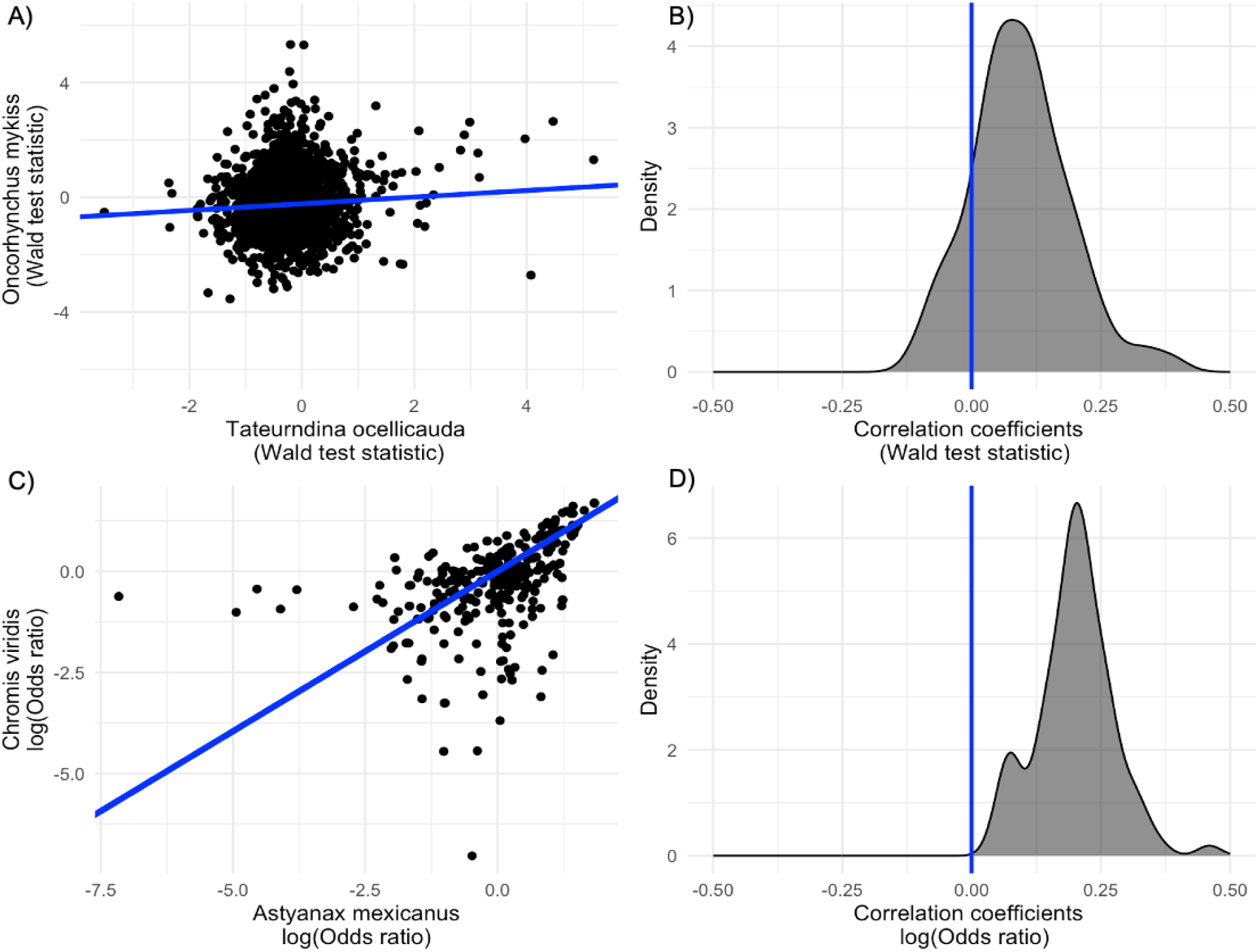
A) Differential orthogroup expression Wald test statistic correlation between *T. ocellicauda* and *O. mykiss* with a linear fit model, and C) gene ontology enrichment odds ratio correlation between *A. mexicanus* and *C. viridis* with the correlation coefficient and linear model inferred intercept. Pairwise species correlation distributions for B) differential orthogroup expression, and D) gene ontology enrichment.

**Figure 4.**
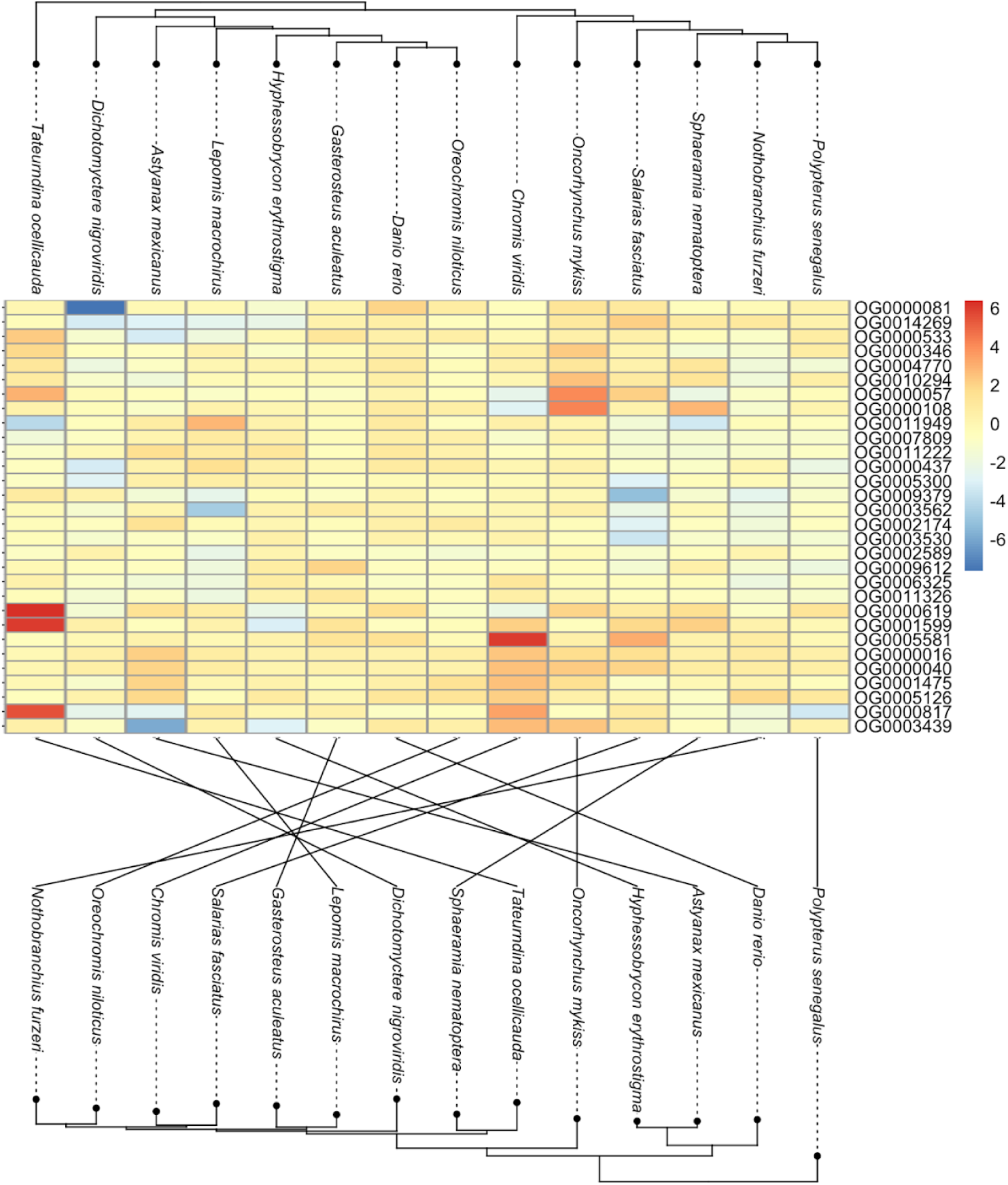
Heatmap representing the top thirty differentially expressed orthogroups with the dendrogram visualizing the hierarchy of expression data (upper), and the species tree inferred from all orthogroups (lower) with lines illustrating the co-phylogeny between dendrogram and species tree.

A similar result is observed for the enrichment of GO terms for differentially expressed transcripts (Wald test, p-value > 0.1) where there are some GO terms which are similarly enriched between species. As an example, *A. mexicanus* and *C. viridis* GO enrichment show a positive correlation (Pearson’s correlation coefficient = 0.33, t = 29.86, p-value < 2.2e-16) (Figure 3C). Overall, species-pairwise correlation estimates for GO enrichment are positive (t-test, t = 26.1, p-value < 2.2e-16) (Figure 3D). Therefore, the response to the immune stimulus evokes specific ontology terms which are utilized in mounting the immune response to the alum treatment.

## Discussion

When comparative biologists observe a phenotype present across most or all members of a clade, they are inclined to conclude that the trait is homologous, and conserved (Wiley & Lieberman 2011). Principles of parsimony (likelihood models of trait evolution) would suggest that the trait evolved prior to the most recent common ancestor of the clade, and has been maintained as the clade diversified. But, does this phenotypic homology imply that the proximate genetic, regulatory, and developmental processes are also homologous? Or, might homology be superficial? Complex biological systems are often characterized by many-to-one mapping rules. That is, there may be many configurations of parts (different bone lengths, cytokine levels, or gene expression levels) that synergistically yield a single outcome. For instance, the nonlinear properties of 4-bar levers imply that many possible fish skull and jaw shapes can yield identical kinematic force transmission for biting or suction feeding (Wainwright et al. 2005). A function (e.g., bite force) could thus be kept constant through evolution (appearing to be homologous), even as its proximal mechanism (skull shape) diverges between species (Alfaro et al. 2005).

The fibrosis immune response is largely conserved across ray-finned fish. Intraperitoneal injection of an immune adjuvant, alum, was recently found to induce fibrosis in threespine stickleback (Hund et al. 2022). Applying this assay to 17 species of Actinopterygian fishes from a wide diversity of sub-clades, Vrtílek and Bolnick (2021) showed that nearly all fishes retain the ability to initiate peritoneal fibrosis (Vrtílek & Bolnick 2021) with the exception of Otophysa species; the Mexican tetra (*A, mexicanus*) and the bleeding-heart tetra (*H. erythrostigma*). This result suggests that the fibrosis immune response (commonly associated with helminth infection) evolved in, or before, the most recent common ancestor of the Actinopterygii, and hence most vertebrates. Fibrosis appears to be a highly conserved homologous trait that has been sustained across more than 350 million years of evolutionary divergence. Our results, however, show that the underlying genetic program is far less homologous.

The conserved inducible fibrosis phenotype provides an opportunity to determine how the associated gene expression response evolves in a phylogenetically informed approach. Specifically, we characterize how associated differential expression is evolving across greater than 350 million years (Hughes et al. 2018; Helfman et al. 2009) of evolutionary divergence present in 14 of the experimental species from Vrtilek and Bolnick (2021). Here, we assigned transcripts to orthogroups which have a shared evolutionary history, revealing the predominant response to the immune stimulus is species-dependent with limited shared treatment effects. Most species do respond to alum injection and most initiate fibrosis (Vrtílek and Bolnick 2021), but each species uses a unique set of expression changes to achieve the apparently homologous phenotypic response. However, orthogroup differential expression was positively correlated across species. Gene ontology enrichment was also positively correlated, which together, suggest each species is evolving independently while maintaining the ability to induce peritoneal fibrosis, yet, there is a degree of functional constraint present with coordinated enriched transcript ontologies. While individual transcripts or orthogroups might be used differently from species to species, across the transcriptome as a whole similar families of genes are nevertheless being engaged, for some (but not all) combinations of species being compared.

As inferred from the differential transcriptomic response to alum, species maintain the ability to induce fibrosis through differing gene regulatory mechanisms, indicating a functional evolutionary constraint, yet, multiple different expression pathways to arrive at the conserved phenotype. This result is contrary to a common assumption in comparative biology. For example, even with a strong selective force like hydrogen sulfide, with documented convergent evolution in morphological and physiological phenotypes in three sympatric fish lineages, the patterns of genomic divergence relative to sulfide tolerance is non-convergent with no consistent differentiated genomic regions (Greenway et al. 2024), The varied way by which fish arrive at the same fibrosis phenotype could be attributed to selection on specific components of the highly complex gene regulatory network which gives rise to fibrosis. Stabilizing selection on the ability to employ fibrosis can reflect its general importance as a physiological function, involved in many biological processes including development, wound healing, and immunity. But this pleiotropy may mean that different species, while maintaining fibrosis, evolve to deploy it in different contexts, prioritizing different subsets of the pleiotropic functions. For instance, the ability to induce fibrosis in response to flatworm infection is heterogeneously maintained in stickleback lake populations separated by only a few kilometers (Weber et al. 2022), facing similar parasite encounter rates. In other species fibrosis may be deployed more in wound-healing, or in embryonic development, etc. These divergent applications of fibrosis may select for divergent gene regulatory pathways, producing the diversity of transcriptomic responses that we observe, even as fibrosis responses are retained. Therefore, even if the phenotypes under selection are conserved, the mechanism by which organisms arrive at the phenotype may not be conserved. As the gene regulatory architecture which gives rise to the fibrosis phenotype is generally inconsistent across the ray-finned fishes, this indicates that evolution is exploring neutral changes in pathways which retain the same function, evoking the concept of systems drift (or developmental systems drift) (True & Haag 2001; Schiffman & Ralph 2022).

Overall, our experimental results represent the incongruence that can arise between the emergence of a conserved immune phenotype (here, fibrosis), and the gene regulatory mechanisms which are induced during the immune response. We demonstrate that vertebrate species responding to a common immune stimulus resulting in the same immune phenotype occurs through species-specific gene regulatory logic. Evaluating the consistency of these results across other vertebrates, particularly those employed in model immune research, highlights a need for phylogenetically diverse immune models systems. Moreover, the diverse gene regulatory structures suggest that no single species will be an effective genetic model for human disease or physiological functions. A diverse portfolio of research organisms will better span the variety of immune regulatory systems in vertebrates.

## Acknowledgments

We gratefully acknowledge the contribution of the Genomic Sequencing and Analysis Facility at University of Texas at Austin. High performance computing resources were provided by the University of Connecticut Computational Biology Core. MV was supported by a Fulbright Commission fellowship for research scholars. The project was funded by the National Institutes of Health (NIAID grant 1R01AI123659-01A1) held by DIB.

## Author contributions

The study was conceived and designed by DIB and MV. MV conducted experimental work. Transcriptome and 3’TagSeq sequencing performed by AR, and LEF. Bioinformatics performed by BAF and LEF. Statistical analyses and graphics by BAF and DIB. Funding acquisition by DIB and MV. Initial draft by BAF and DIB with comments and feedback from LEF, AR, and MV.

## Supplemental Figures

**Figure S1:**
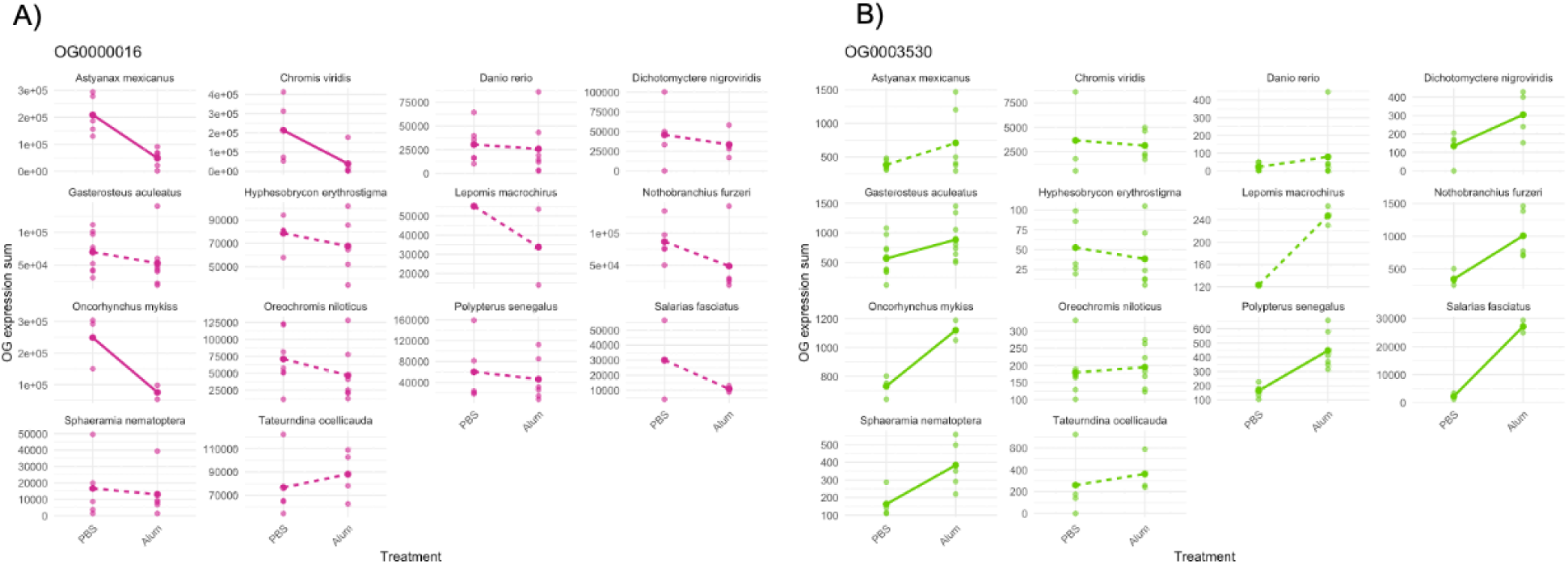
Orthogroup expression for PBS and Alum injections across all species where solid lines represent a significant treatment effect (p-value < 0.1) and dotted line represents non-significant treatment effect (p-value > 0.1) for A) orthogroup with highest effect size of treatment (OG0000016), and B) orthogroup with highest effect size of the interaction of treatment and species (OG0003530).

**Figure S2:**
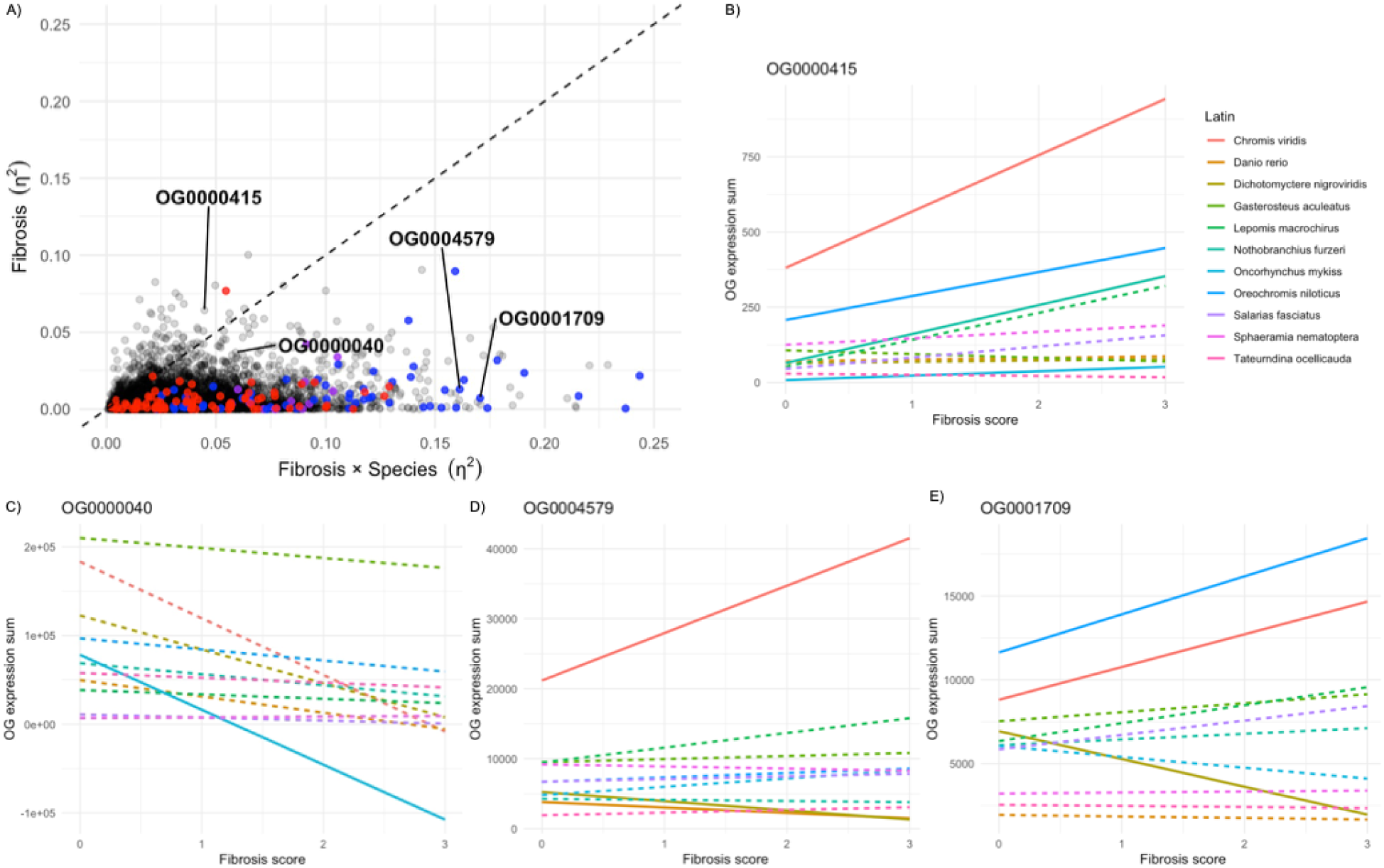
A) For each orthogroup summed normalized gene expression, we used a linear model to estimate the effect size (η^2^) for the effects of fibrosis score, and the fibrosis score × species interaction on variation in the orthogroup expression. Treatment η^2^ indicates the extent to which any orthogroup summed normalized expression diverges predictably for fibrosis. The η^2^ for the fibrosis × species interaction measures the extent to which fibrosis depends on the species for every orthogroup. The dashed line is a 1:1 line, for ease of visualization; points falling above this line have a larger fibrosis effect than interaction effect with blue significant effect of fibrosis only, purple fibrosis and interaction, and red only the interaction term (p-value < 0.1). Orthogroup summed expression for B) OG0000415, C) OG0000040, D) OG0004579), and E) OG0001709 where solid lines represent a significant fibrosis effect (p-value < 0.1).

## Notes

### Competing Interest Statement

The authors have declared no competing interest.

